# Lysine-independent ubiquitination and degradation of REV-ERBα involves a bi-functional degradation control sequence at its N-terminus

**DOI:** 10.1101/2023.05.01.538963

**Authors:** Ting-Chung Suen, Jason P. DeBruyne

## Abstract

REV-ERBα and REV-ERBβ proteins play crucial roles in linking the circadian system to overt daily rhythms in mammalian physiology and behavior. In most tissues, REV-ERBα protein robustly cycles such that it is detected only within a tight interval of 4-6 hours each day, suggesting both its synthesis and degradation are tightly controlled. Several ubiquitin ligases are known to drive REV-ERBα degradation, but how they interact with REV-ERBα and which lysine residues they ubiquitinate to promote degradation are unknown. In this study, we attempted to identify both ubiquitin-ligase-binding and ubiquitination sites within REV-ERBα required for its degradation. Surprisingly, mutating all lysine residues, the common sites for ubiquitin conjugation, in REV-ERBα to arginines (K20R), did very little to impair its degradation in cells. K20R were degraded much faster by co-expression of two E3 ligases, SIAH2 or SPSB4, suggesting possible N-terminal ubiquitination. To explore this, we examined if small deletions at the N-terminus of REV-ERBα would alter its degradation. Interestingly, deletion of amino acid (AA) residues 2 to 9 (delAA2-9) clearly resulted in a less stable REV-ERBα. We found that it was the length (i.e. 8 AA), and not the specific sequence, that confers stability in this region. Simultaneously, we also mapped the interaction site of the E3 ligase SPSB4 to this same region, specifically requiring AA4-9 of REV-ERBα. Thus, the first 9 AA of REV-ERBα has two opposing roles in regulating REV-ERBα turnover. Further, deleting eight additional AAs (delAA2-17) from the N-terminus strongly prevents REV-ERBα degradation. Combined, these results suggest that complex interactions within the first 25AAs potentially act as an endogenous ‘switch’ that allows REV-ERBα to exist in a stabilized conformation in order to accumulate at one time of day, but then rapidly shifts to a destabilized form, to enhance its removal at the end of its daily cycle.

## Introduction

Proper functioning of the circadian clock depends on timely expression of multiple transcription factors that comprise a set of interlocking transcriptional-translational negative feedback loops (1–3). REV-ERBα and REV-ERBβ are transcriptional repressors that are important components of the mammalian circadian clock (4–7). Expression of *Rev-Erbα* at the RNA level is robustly rhythmic, and REV-ERBα protein is only detectable within a 4-hour interval during the day (4). The tight interval of REV-ERBα protein existence in the cells highlights the importance of its rapid elimination from the cells at the appropriate time. However, the mechanisms of how such fast degradation being accomplished at the appropriate time is unknown.

Most proteins in the cells are eliminated by the ubiquitin-proteasome (UPS) system, where chains of ubiquitin molecules are covalently attached to one or more lysine residues on proteins targeted for degradation. This important first step of protein degradation is achieved through the concerted actions of a cascade of enzymes, E1 ubiquitin activating enzymes, E2 ubiquitin conjugating enzymes, and E3 ubiquitin ligases (8–10). Ubiquitin ligases provide specificity on which proteins are to be targeted and degraded by the proteasome (11–14). Our laboratory developed a functional screen (15) that could identify which ubiquitin ligase(s) is responsible for degrading a specific protein of interest. SIAH2, SPSB1, and SPSB4 were identified in this screening strategy to strongly accelerate the degradation of REV-ERBα (15). Depletion of SIAH2, SPSB1, and SPSB4 resulted in the alteration of REV-ERBα level and suppression of its downstream target genes, as well as the overall circadian oscillator function (15, 16), suggesting that timely degradation of REV-ERB is important for proper clock function. Interestingly, study on SIAH2 knockout mice revealed itself as a female-specific regulator of circadian rhythms and metabolism, although the role of REV-ERBα in this was not clear (17). Other ubiquitin ligases, Arf-bp1 and Pam (18), FBXW7 (19), have also been reported to drive degradation of REV-ERBα. Removal of each of these E3 ligases have distinct effects on REV-ERBα stability and overall clock function, suggesting that each serves a specific, non-overlapping role in REV-ERBα degradation. In this study, we focused on identifying the basic regulatory mechanisms intrinsic to the REV-ERBα protein itself, as a starting point for determining how the different E3 ligases may cooperate. To do this, we began with genetically mapping E3 binding and ubiquitination sites on the REV-ERBα protein. We found however, that lysine-mediated ubiquitination is not required for REV-ERBα degradation, and instead may utilize an intrinsic bifunctional “switch” located at its N-terminnus to both provide stability and facilitate its degradation by the proteasome. This novel dual-functionaing domain potentially could control the timing of a transition of REV-ERBα from an accumulation state to an accelerated degradation state, contributing to its overall daily cycling in abundance and function.

## Materials and Methods

### Cell lines and transfection

The human embryonic kidney cell line HEK293 was maintained in Dulbecco’s modified Eagle’s medium (DMEM) (Corning Cat. No. 10-017-CV) with 10% fetal bovine serum (VWR Cat. No. 89510-186), in a 37^0^C humidified incubator with 5% CO2. Transfection was carried out with Fugene HD (Promega, Cat. No. E2311, Madison, WI) or Invitrogen Lipofectamine 2000 Transfection Reagent (Cat. No. 11668500, Fisher Scientific, Waltham, MA) according to manufacturer’s instructions.

### Plasmids and mutants

pCMVSport6Rev-Erbα contains the mouse *rev-erbα* cDNA cloned into the mammalian expression vector pCMVSport6. *In vitro* mutagenesis was carried out with the Q5 site-directed mutagenesis kit (Cat. No. E0554S, New England Biolabs, Ipswich, MA). Primers specific for the desired mutations were designed with the NEBaseChanger on the NEB website, and were purchased from Thermo Fisher Scientific (Waltham, MA). Briefly, 10ng of plasmid was mixed with 1mM of a specific primer pair, along with the enzyme mix provided. PCR was carried out as described in the mutagenesis protocol with annealing temperature selected for the specific primer pair. PCR products were then checked by agarose gel electrophoresis. Products with the correct size were treated with a mixture of enzymes including kinase, ligase, and Dpnl (KLD mix) for 10 minutes, such that the original plasmid would be digested to reduce background during bacterial transformation. 1ml of the KLD-treated newly synthesized plasmid was used to transform the DH5a strain of *E. coli* provided. Colonies were picked and grown overnight in Luria Broth medium with appropriate antibiotics, plasmids were isolated with Qiagen (Germantown, MD) Plasmid Mini Kit (Cat. No. 12125), and DNA were sent for sequencing with appropriate primer to confirm if the specific mutation was successfully introduced.

### Primers

Oligonucleotides or primers were purchased from Thermo Fisher Scientific (Waltham, MA). Information for all primers will be available upon request.

### Sequencing

All mutants used in this study were confirmed by sequencing, performed either by the Core Facility of Morehouse School of Medicine (MSM) or Eurofins Genomics (Louisville, KY), with sequencing primer chosen specifically to verify if the intended mutations were successfully obtained.

### Antibodies

Antibodies were obtained from Cell Signaling Technology (Danvers, MA). These include anti-REV-ERBα (Cat No. 13418S), anti-GAPDH (Cat. No. 5174S), anti-β-Actin (Cat. No. 4970S), anti-HA-Tag (rabbit, Cat. No. 3724S, and mouse, Cat. No. 2367S), anti-MYC-tag (rabbit, Cat. No. 2278S, and mouse, Cat. No. 2276S), HRP-conjugated secondary antibodies include Goat anti-mouse IgG (Cat. No. 91196) and anti-rabbit IgG (Cat. No. 7074). Fluorescence conjugated secondary antibodies were purchased from Fisher Scientific (Waltham, MA), these include Invitrogen goat-anti-rabbit (Cat. No. PIA32735) or goat-anti-mouse (Cat. No. PIA32730) Alexa Fluor Plus 800, goat-anti-rabbit (Cat. No. PIA32734) or goat-anti-mouse (Cat No. PIA32729) Fluor Plus 680. Protein-A agarose beads (Cat No. 9863S) were also products of Cell Signaling Technology.

### Protein degradation assays

Typically, HEK293 cells were washed with Phosphate Buffered Saline (PBS, Cytiva HyClone Cat. No. BSS-PBS-1X6), trypsinized (Corning 0.25% Trypsin, Cat. No. 25-053-CI), collected with DMEM, and centrifuged (1500 rpm, 5 minute). Cells were resuspended in fresh medium, counted, and seeded into a 24-well plate (Corning Falcon 353047) at a density of 10^5^ cells per well. 100 ng of plasmids that express the cDNA of either the wildtype (WT) or mutant REV-ERBα were transfected into multiple wells with Fugene HD (Promega, Cat. No. E2311, Madison, WI) or Invitrogen Lipofectamine 2000 Transfection Reagent (Cat. No. 11668500, Fisher Scientific). For experiments on ubiquitin ligase-induced degradation, 50 ng of empty vector pCMVSport6, SIAH2-, or SPSB4-expressing plasmid, was added to the 100ng of REV-ERBα expressing plasmid. 36-48 hours after transfection, cells were treated with cycloheximide (Cat. No. 2112S, Cell Signaling, Danvers, MA) at 10 μg/ml final concentration for various times as indicated in individual experiments. The proteasome inhibitor MG132 (Cat. No. 2194S, Cell Signaling, Danvers, MA) was also added to one of the wells at a final concentration of 5μM, usually treated along with the longest time point of cycloheximide. Cells were washed with PBS, followed by lysis with 100μl of RIPA buffer (diluted from a 2X solution, Cat. No. BP115X, Boston Bioproducts Inc., Ashland, MA), supplemented with a protease inhibitor cocktail (Cat. No. 5871S, Cell Signaling, Danvers, MA), at 4^0^C on a rocking platform. Cell lysates were collected after one to two hour and transferred from the individual wells into microcentrifuge tubes. Cell debris were cleared by centrifugation at 13,000 rpm in a microcentrifuge at 4^0^C for 15-30 minutes. Supernatants were then transferred to new tubes and either stored at -80^0^C, or were ready for Western analysis.

### Western blots

Aliquots of cell lysates were mixed with appropriate amount of a 6X SDS-Sample buffer, reducing (Cat. No. BP-11R, Boston Bioproducts Inc., Ashland, MA), heated to 95^0^C for 5 minutes, before being loaded onto a Bio-Rad Criterion TGX or mini-PROTEAN TGX gel, and run with the Criterion Vertical Electrophoresis Cell system (Bio-Rad, Hercules, CA). Proteins were then transferred to Cytiva Amersham Hybond PVDF membranes (formerly GE Healthcare Life Sciences, now Global Life Sciences Solution USA LLC, Marlborough, MA) using the Bio-Rad Trans-Blot Turbo transfer system. Membranes were washed for 5 minutes at room temperature on an orbital shaker with 1X Tris-buffered saline (TBS), diluted from a 20X TBS stock solution (Cat. No. BM300X, Boston Bioproducts Inc., Ashland, MA).

Blocking was followed by placing the membrane in 1X TBS with 5% non-fat dry milk or bovine serum albumin (BSA) (Boston Bioproducts Inc., Ashland, MA), on a rocking platform at 4^0^C. After an hour, an appropriate antibody in TBST (TBS with 0.1% Tween-20) with 5% of either non-fat dry milk or BSA was then incubated with the membrane on a rocking platform at 4^0^C. After overnight incubation, membranes were washed with 1X TBST for 10 minutes at room temperature on an orbital shaker. After 3 washes, appropriate secondary antibody in TBST was added and left shaking for 1 hour at room temperature on the rotating platform. Three washes with TBST were then carried out as before.

Membranes were exposed to chemiluminescence when HRP-conjugated antibody was used, while imaging was obtained with LI-COR Odyssey Fc Imaging system (LI-COR, Lincoln, NB) when fluorescein-conjugated antibodies were used.

### Immunoprecipitation

Typically, 500 ng of the plasmid vector pCMVSport6, pCMVSport6REV-ERBα-WT or mutants were transfected into different wells of a 6-well tissue culture plate (Corning Falcon 353046), with either 250ng of pCMVSport6 or pCMVSport6-HA-Ub, a plasmid that expresses a HA-tagged ubiquitin. 36-48 hours after transfection, cells were changed into medium containing 5 μM MG132 (Cat. No. 2194S, Cell Signaling, Danvers, MA). 2-3 hours later, cells were washed with 2-ml PBS followed by the addition of 300μl trypsin. Detached cells were collected into 15 ml conical tubes with 4ml of medium. Cells were centrifuged at room temperature for 5 minutes at 1,500rpm. The supernatants were removed, and the cell pellets were resuspended with 1.5ml PBS and transferred to microcentrifuge tubes. Cells were centrifuged at 1,500rpm for 5 minutes in 4^0^C, PBS was aspirated off and the pellets were resuspended with 150μl of RIPA buffer with protease inhibitors. The tubes were then left on a rocking platform at 4^0^C for 1-2 hours. Cell lysates were then cleared by spinning at 13,000rpm for 30 minutes at 4^0^C, after which a 10 μl aliquot was transferred to a tube with 2 μl of Laemmli 6X SDS-Sample buffer and stored as input lysate in -80^0^C. The remaining cell lysates were added to Eppendorf tubes containing 650 μl of PBST (phosphate buffered saline with 0.1% Tween-20) with protease inhibitor cocktail, along with 10 μl of pre-washed Protein-A Agarose beads (Cat. No. 9863S, Cell Signaling, Danvers, MA). The tubes were left on a rotating disc at 4^0^C overnight for pre-clearing the lysates. The agarose beads were spun down at 4,000rpm for 3 minutes in a microcentrifuge at 4^0^C. Cell lystaes were then transferred to low-protein binding Eppendorf tubes (Cat. No. 13-698-794, Fisher Scientific) containing 3 μl of anti-REV-ERBα antibody. Tubes were placed back to the rotating disc at 4^0^C for 4-8 hours, after which a fresh 15 μl of Protein-A Agarose beads were added. The tubes were again placed back onto the rotating disc overnight. The immunoprecipitated mix was centrifuged at 4,000 rpm for 3 minutes, supernatant removed, and fresh PBST was added to wash the agarose beads with the tubes back onto the rotating disc. Centrifugation and washing were repeated 3 more times. Supernatant was carefully removed as much as possible after the last wash. 25 μl of 6X SDS-Sample buffer, reducing loading dye, were added to the beads. The immunoprecipitated mix, along with the input lysates aliquoted the day before were heated to 95^0^C for 10 minutes, vortexed briefly and spun down in microcentrifuge, before being loaded on a gel for Western.

## Results

### Lysine-independent degradation of REV-ERBα

Since the first step to mark a protein for degradation is the addition of a chain of ubiquitin molecules to one or more lysine residues on the protein (8–10), our initial approach was to try to identify which lysine residues are important for the degradation of REV-ERBα. A simple schematic illustration of the mouse REV-ERBα protein is shown in Figure 1A, with the DNA-binding and ligand-binding domains shown as grey and open boxes, respectively. The relative positions of all lysine residues are indicated by “K”s (not drawn to scale). Three ubiquitinated peptides were found in an ubiquitinome study (20) and mapped to lysines at amino acid (AA) positions 456, 577, and 592 (the three underlined “K”s in Figure 1A) on REV-ERBα. These 3 lysines were therefore mutated to arginines (R), both individually and in combinations, to determine if they are required for REV-ERBα’s degradation. For these experiments, we used a HEK293 cell-based degradation assay that allowed us to both assess REV-ERBα stability in the cells, with or without co-expressing E3 ligases. When these mutants were transfected into HEK293 cells, their degradation profiles in a cycloheximide time course were very comparable to the wildtype (WT) protein, suggesting that these three lysines are not required for REV-ERBα degradation (data not shown). After multiple rounds of mutating additional lysine and stability testings, we ended up with a REV-ERBα protein with all its twenty lysines changed to arginines, (the K20R mutant in Figure 1A). To our surprise, although removing all lysine residues slowed down the degradation of REV-ERBα (Figure 1B, compare lanes 9-15 to lanes 1-7), the mutant protein was still efficiently degraded by the end of the time course (Figure 1C). The degradation was clearly mediated by proteasome as the level of REV-ERBα was maintained after 6 hours when the proteasome inhibitor MG132 (lanes ‘M’) was added to the cells (lanes 8 and 16 compared with lanes 1 and 9, respectively). Thus, although lysine ubiquitination is known to be the major and essential initiation event that leads to protein degradation by the proteasome (17), the above results clearly showed that ubiquitination on lysine residue is not a requiremnet for proteasome-mediated degradation of REV-ERBα.

**Figure 1.**
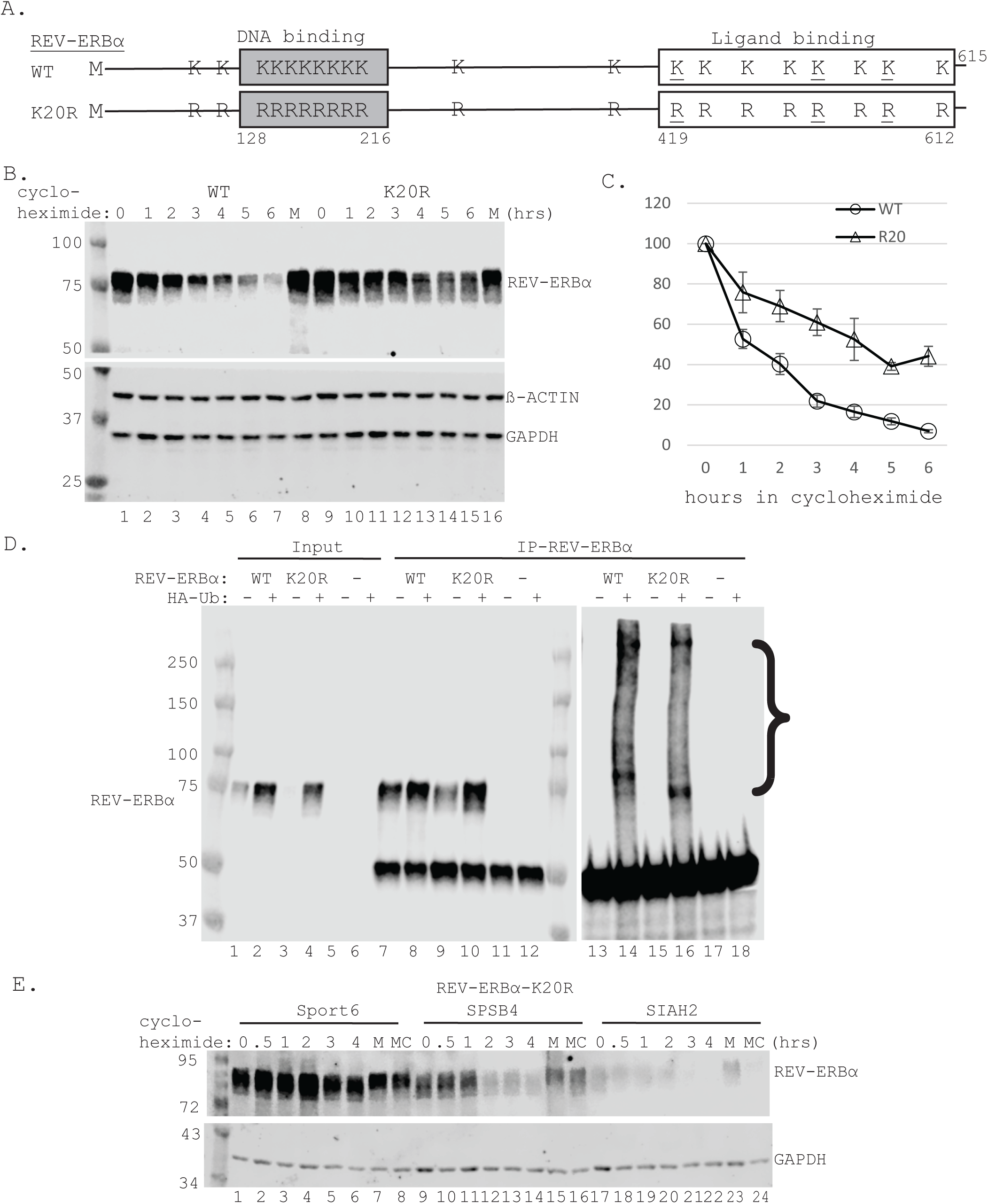
Lysine-independent ubiquitination and degradation of REV-ERBα. A. Wildtype (WT) mouse REV-ERBα is 615 amino acids (AAs) in length, where methionine (M) is at position 1. The 20 lysines (K) are distributed with respect to the DNA-binding (grey box, AA128-216) and ligand-binding (open box, AA419-612) domains, not drawn to scale. The 3 underlined Ks were shown to be ubiquitinated and published in an ubiquitinome study. All Ks are changed to Arginines (Rs) by *in vitro* mutagenesis and confirmed by sequencing in the K20R mutant. B. WT (lanes 1-8) or K20R (lanes 9-16) were transfected into HEK293 cells seeded in 24-well plate. 36-48 hours after transfection, cells were treated with cycloheximide for the indicated number of hours, and the Western blot was probed with anti-REV-ERBα antibody. The membrane was then probed with anti-glyceraldehyde-3-phosphate-dehydrogenase (GAPDH) antibody, and anti-βACTIN antibody, as loading controls. Lanes “M” were lysates from cells treated with MG132 for 6 hours (lanes 8 and 16). C. Intensity of REV-ERBα and GAPDH bands were measured with the Li-COR ImageStudio software. Quantities of the WT and R20 were normalized with GAPDH and all time points were expressed as a relative number to the respective time ‘0”, which is set at 100. Results are the means of three experiments. D. WT or K20R were transfected into HEK293 cells with (“+” lanes) or without (“-” lanes) co-transfection of a plasmid that expresses a HA-tagged ubiquitin (HA-Ub). Lysates from transfected cells (lanes 1-6) were immunoprecipitated (IP) with an anti-REV-ERBα antibody (lanes 7-18). Western blots were probed with an anti-REV-ERBα (lanes 1-12) or anti-HA antibody (lanes 13-18). Bands that form a ladder are marked with a bracket on the right, representing REV-ERBα proteins that have increasing number of ubiquitin attached. E. K20R were co-transfected with Sport6 (lanes 1-8), Sport6-SPSB4 (lanes 9-16), or Sport6SIAH2 (lanes 17-24), into HEK293 cells. 36-48 hours after transfection, cells were treated with cycloheximide for the indicated number of hours, and the Western blot was probed with anti-REV-ERBα antibody. Lanes “M” were lysates from cells treated with MG132 for 4 hours (lanes 7, 15, and 23). Lanes “MC” were lysates from cells treated with both MG132 and cycloheximide for 4 hours (lanes 8, 16, and 24). Anti-GAPDH was used as a loading control.

### Lysine-independent ubiquitination of REV-ERBα

The delayed but efficient degradation of K20R prompted us to test if it could still be ubiquitinated when compared to the WT protein under identical conditions. Both WT and K20R were expressed in HEK293 cells (Figure 1D, “Input” lanes 1-4), with (“+” lanes) or without (“-” lanes) co-transfection of a plasmid that expresses a HA-tagged ubiquitin cDNA (HA-Ub). Cells were lysed and followed by immunoprecipitation (IP) with anti-REV-ERBα antibody (lanes 7-18). Western blot using anti-REV-ERBα antibody (lanes 1-12) showed that similar amount of WT and K20R were successfully brought down by IP (lanes 7-10), while endogenous REV-ERBα was undetectable in non-transfected HEK293 cells (lanes 5, 6, 11, 12). When anti-HA antibody was used (lanes 13-18), bands appearing as ladders (marked by bracket) were detected only in those lanes where HA-Ub were also co-transfected (lanes 14 and 16). The absence of these ladders in the HEK293 cells (lanes 17 and 18) further confirmed their specificity. These bands all have higher molecular weight than that of REV-ERBα, suggesting that they are REV-ERBα with increasing numbers of ubiquitin molecules attached, as each additional ubiquitin molecule would increase the size of the protein by 8kD. Importantly, the ladder of bands detected in K20R (lane 16) suggested that it is also efficiently ubiquitinated even though it does not even have a single lysine. This result was intriguing as the ε-amino group on lysine is known to be the site where ubiquitin attaches to proteins. Nevertheless, the ability of K20R to be ubiquitinated is consistent with the fact that it can be readily targeted to the proteasome for degradation (Figures 1B and 1C).

Our laboratory previously identified several ubiquitin ligases, SIAH2, SPSB1, and SPSB4 (15, 16), that were able to accelerate the degradation of REV-ERBα when they were co-expressed. Depletion of these E3 ligases also resulted the suppression of target genes downstream of REV-ERBα, as well as an extended circadian period, (15, 16). It is commonly accepted that E3 ligases function to ubiquitinate proteins on lysine residue(s), we therefore tested if E3-induced degradation of REV-ERBα was affected in the K20R mutant. Consistent with results in Figures 1B and 1C, the K20R mutant is relatively stable in the first 4 hours (Figure 1E, lanes 1-8). When SPSB4 was co-expressed, K20R degradation was clearly accelerated (lanes 9-16). SIAH2’s ability to degrade K20R appeared to be much more potent than SPSB4, as only weak signals of K20R were detected (lanes 17-24). These data further support the noton that ubiquitination and proteasomal degradation of REV-ERBα may be lysine-independent.

### N-terminus of REV-ERBα controls its degradation

The ε-amino group of lysine residue is commonly reported as the predominant ubiquitin attachment site driven by ubiquitin ligases (8–10), a likely site of ubiquitin conjugation on the K20R mutant would therefore be the only free α-amino group on the methionine located at the very N-terminus of the protein, as reported in a handful of proteins (21–25). We reasoned that if the N-terminus is important for ubiquitination, perhaps altering the amino acids in its proximity would affect degradation. Since the N-terminal methionine is the translation initiation site that needs to be preserved, we introduced small deletions of 8 and16AAs near the N-terminus, (Figure 2A). As shown in the western blot (Figure 2B), the mutant delAA2-9 protein (lanes 9-15) clearly degraded faster than the WT (lanes 1-7, and densitometry quantification shown in Figure 2C). The degradation of delAA2-9 mutant, like the WT protein, appears to be mediated by the proteasome, since the level of protein remained after 10 hours in the presence of MG132 was close to the level found at time 0 (compare lane 8 to lane 1 for WT, lane 16 to lane 9 for delAA2-9). In contrast, extending the deletion to 16AAs (mutant delAA2-17) resulted in a highly stable protein (lanes 17-23, and Figure 2C), one that appeared more stable than the lysine-less K20R mutant (Figure 1B). Interestingly, the presence of MG132 was not able to rescue the level of delAA2-17 mutant back to the level found at time 0 (compare lane 24 to lane 17). Instead, the level of delAA2-17 found after 10 hours of MG132 treatment (lane 24) was very similar to the level found after 10 hours of cycloheximide treatment (lane 23), suggesting that its slow degradation was not mediated by proteasome.

**Figure 2.**
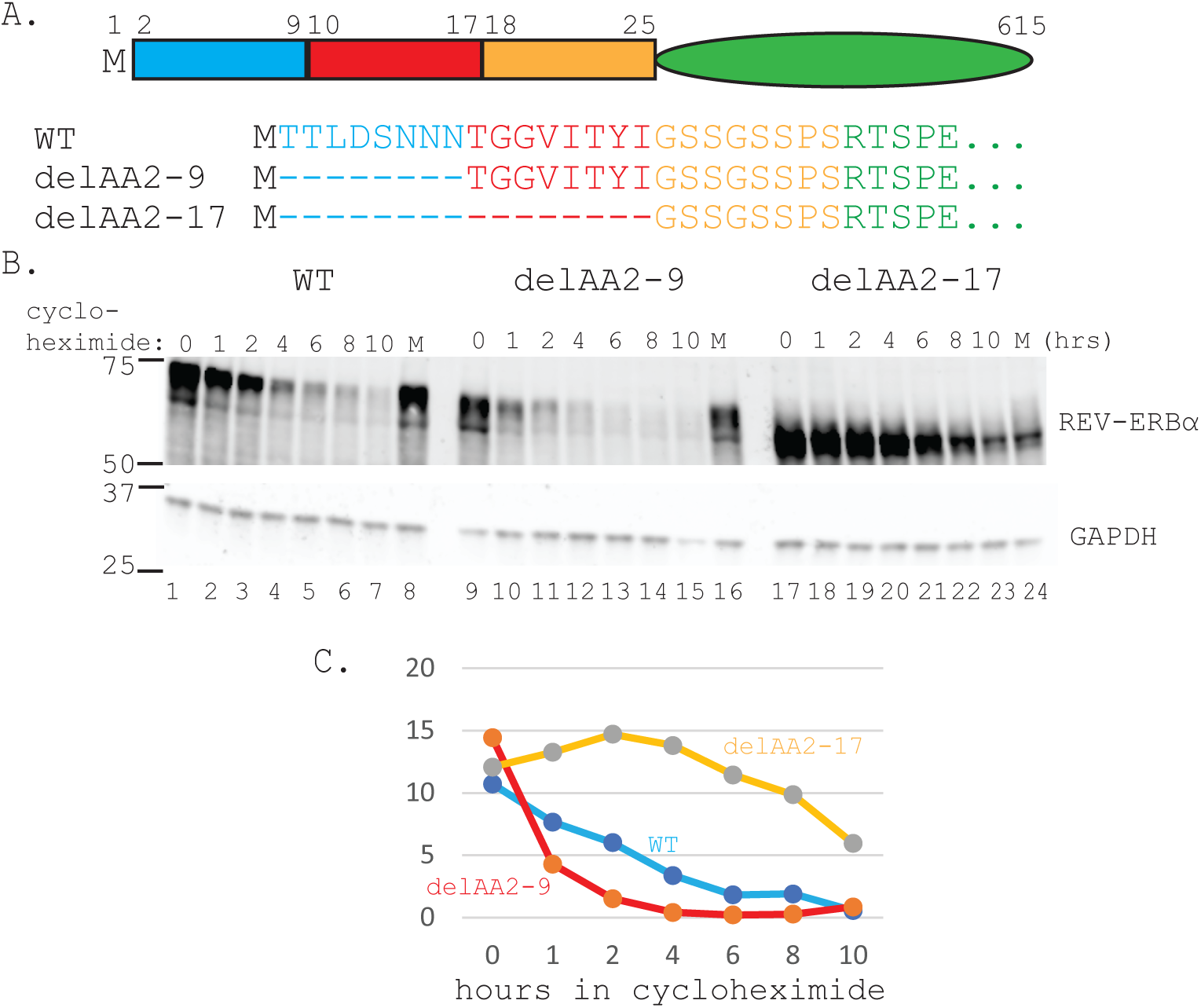
Small deletions at the N-terminus alters REV-ERBα stability. A. Wildtype (WT) REV-ERBα is drawn not to scale, with the first 25 amino acids (AAs) enlarged as three colored boxes, each 8AAs in length, after the methionine (M) at position 1, the rest of the protein is shown as a green ellipse. The sequence of the first 30AAs is shown underneath, color-matched to the 3 boxes, with dashed lines representing the deletions. B. WT (lanes 1-8) or the indicated mutants (lanes 9-24) were transfected into HEK293 cells seeded in 24-well plate. 36-48 hours after transfection, cells were treated with cycloheximide for the indicated number of hours, and the Western blot was probed with anti-REV-ERBα antibody. The membrane was then probed with anti-glyceraldehyde-3-phosphate-dehydrogenase (GAPDH) antibody, as a loading control. Lanes “M” were lysates from cells treated with MG132 for 10 hours (lanes 8, 16, and 24). C. Intensity of REV-ERBα and GAPDH bands were measured with ImageQuant LAS4000 software provided with the Luminescent Image Analyzer (FujiFim Medical System USA). Arbitrary units were used directly from the software.

Overall, the combined results indicated that the N-terminal 17AA of REV-ERBα contains two small domains that play opposing roles in regulating its degradation.

### Characterization of how AA2-9 may prevent fast degradation REV-ERBα

The removal of AA2-9 from the N-terminus of REV-ERBα clearly resulted in an unstable protein (delAA2-9, Figure 2), suggesting that AA2-9 may have a role in slowing down degradation when the protein is in its native conformation. To examine if there is a sequence requirement for AA2-9 to impede degradation, successive pairs of AA2-3, 4-5, 6-7, or 8-9, respectively, were mutated to double alanines (Figure 3A) and their stability were tested. As shown in Figures 3B and 3C, no discernable difference could be seen with all these substitution mutants when compared to the wild type (lanes 1-6). It is possible that mutating only 2 out of 8AA in this region is not sufficient to disrupt the stabilizing mechanism. We therefore examined two additional mutants, both of which maintained the same AA2-9 composition but they were rearranged in two different ways (Figure 3D). As shown in Figure 3E, reversing the native order of AA2-9 to AA9-2 (revAA9-2, lanes 7-12) did not change the rate of degradation. The other mutant with the order of AA2-9 scrambled (SCRAM-AA2-9, lanes 13-18) appeard to cause a slight destabilization compared to the WT protein (lanes 1-6), but still far more stable than when AA2-9 is deleted (Figure 2). Combined results form this series of mutants suggested that the stability provided by AA2-9 is not mediated by a consensus sequence, such as a specific protein-binding site. Since we performed these experiments to explore the possibility of N-terminal ubiquitination, we decided to examine if a minimal length of N-terminus is needed to cause protein stability. Interestingly, deleting two (delAA2-3, lanes 7-12) or six amino acids (delAA2-7, lanes 13-18) in this region appeared to cause slight but noticeable instability as they were degraded somewhat faster than the WT protein (lanes 1-6, Figure 3G). However, neither mutants caused instability comparable to the highly unstable delAA2-9 (lanes 19-24, Figure 3G). These results suggested that a minimum of two amino acids (N8N9) are enough to delay protein degradation, but AA2-9 is required to provide robust stability. The difference in stability conferred by only 2AA is surprising, and combined with data from above for the specific AA sequence (in particular the N8N9AA mutant), suggested that the length, not the specific sequence is an important factor contributing to the stability.

**Figure 3.**
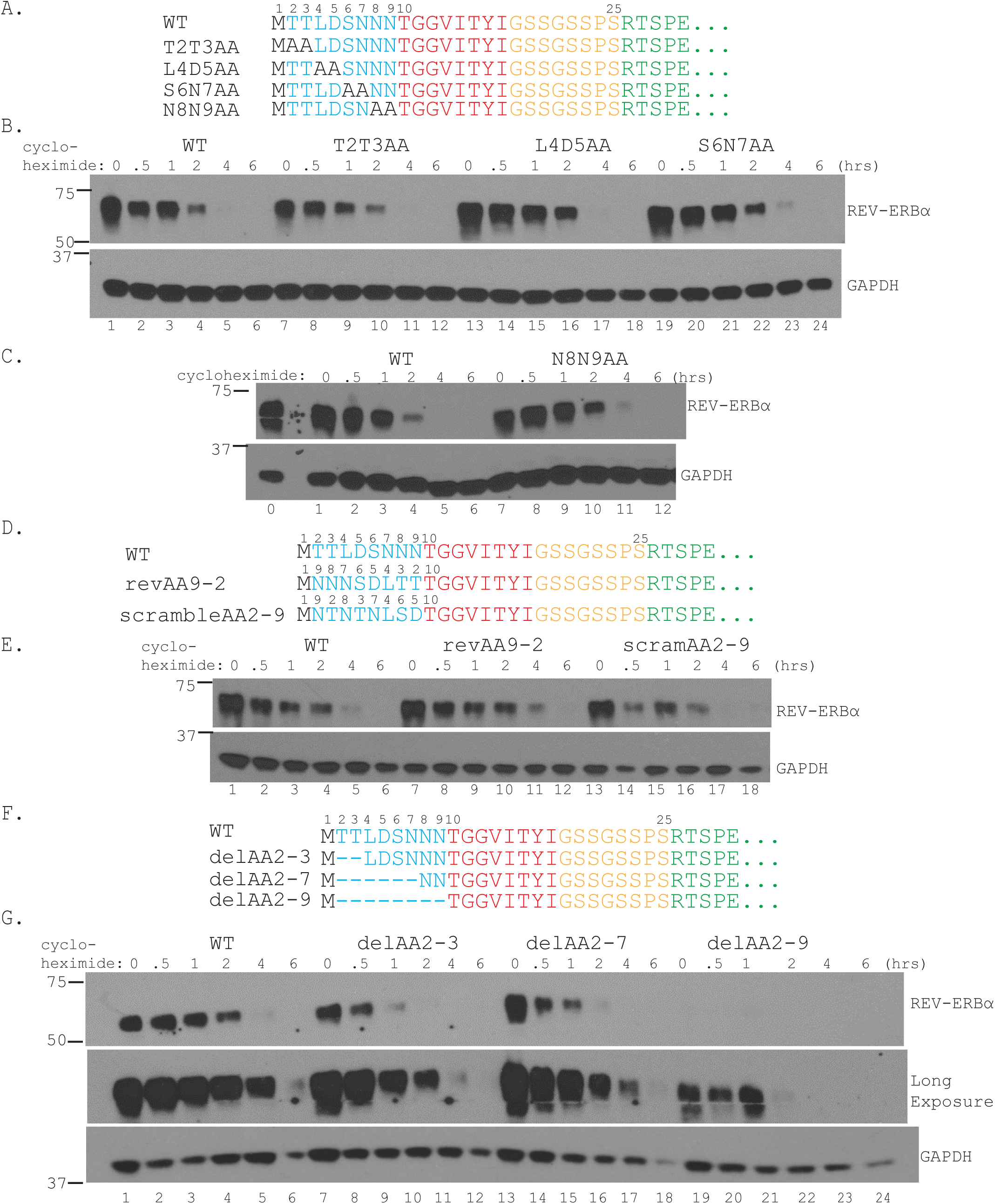
AA2-9 protects REV-ERBα against fast degradation. A. Only the first 30AAs of REV-ERBα is shown as in Figure 2, with the green sequence starting at AA26 representing the rest of the protein. AAs 2-3, 4-5, 6-7, or 8-9, were substituted with double alanine to generate the four mutants as shown. B. and C. WT (lanes 1-6) or the indicated mutants were transfected into HEK293 cells seeded in 24-well plate. 36-48 hours after transfection, cells were treated with cycloheximide for the indicated number of hours, and the Western blot was probed with anti-REV-ERBα antibody. The membrane was then probed with anti-glyceraldehyde-3-phosphate-dehydrogenase (GAPDH) antibody, as a loading control. The WT sample at time “0” (Figure 3B, lane 1) was also loaded into the gel in Figure 3C as lane “0” for cross-membrane comparison. D. AA2-9 sequence was numbered on top as such in the WT and re-arranged as shown in the two mutants. E. WT or mutants in Figure 3D were transfected into HEK293 cells seeded in 24-well plate. 36-48 hours after transfection, cells were treated with cycloheximide for the indicated number of hours, and the Western blot was probed with anti-REV-ERBα antibody. The membrane was then probed with anti-glyceraldehyde-3-phosphate-dehydrogenase (GAPDH) antibody, as a loading control. F. Small deletions in AA2-9 were made as shown. G. WT or mutants in Figure 3F were transfected into HEK293 cells seeded in 24-well plate. 36-48 hours after transfection, cells were treated with cycloheximide for the indicated number of hours, and the Western blot was probed with anti-REV-ERBα antibody. The membrane was then probed with anti-glyceraldehyde-3-phosphate-dehydrogenase (GAPDH) antibody, as a loading control. A long exposure shown at the bottom was needed to detect delAA2-9 (lanes 19-24).

### AA2-9 must be at the very N-terminus to exert its protective effect against degradation

If the presence of AA2-9 at the N-terminus is the reason why REV-ERBα degradation is maintained at the WT level, then mechanistically, it may be hidden or exposed via conformational change as part of the natural degradation process. We explored this possibility by moving AA2-9 to position as AA18-25, and examined the stability of this re-organized mutant (AA2-9insAA25 in Figure 4A). Interestingly, the stability of AA2-9insAA25 (Figure 3B, lanes 19-24) was much more comparable to the delAA2-9 mutant (lanes 7-12), both much less stable than either the WT (lanes 1-6) or the reversed mutant proteins (revAA9-2, lanes 13-18). These results suggest that although AA2-9 is near its native location, it can no longer protect the protein from degrading like the WT. These data suggest that AA2-9 must reside at the very N-terminus in order for REV-ERBα to maintain WT-like stability. Altogether, our data to this point are consistent with the notion that, a conformational change within the N-terminus may in some ways incapacitate AA2-9’s degradation-impeding function. These results also lends support to the hypothesis that possible interactions within the first 25 amino acids of REV-ERBα could indeed function as a switch to alter the speed of degradation.

**Figure 4.**
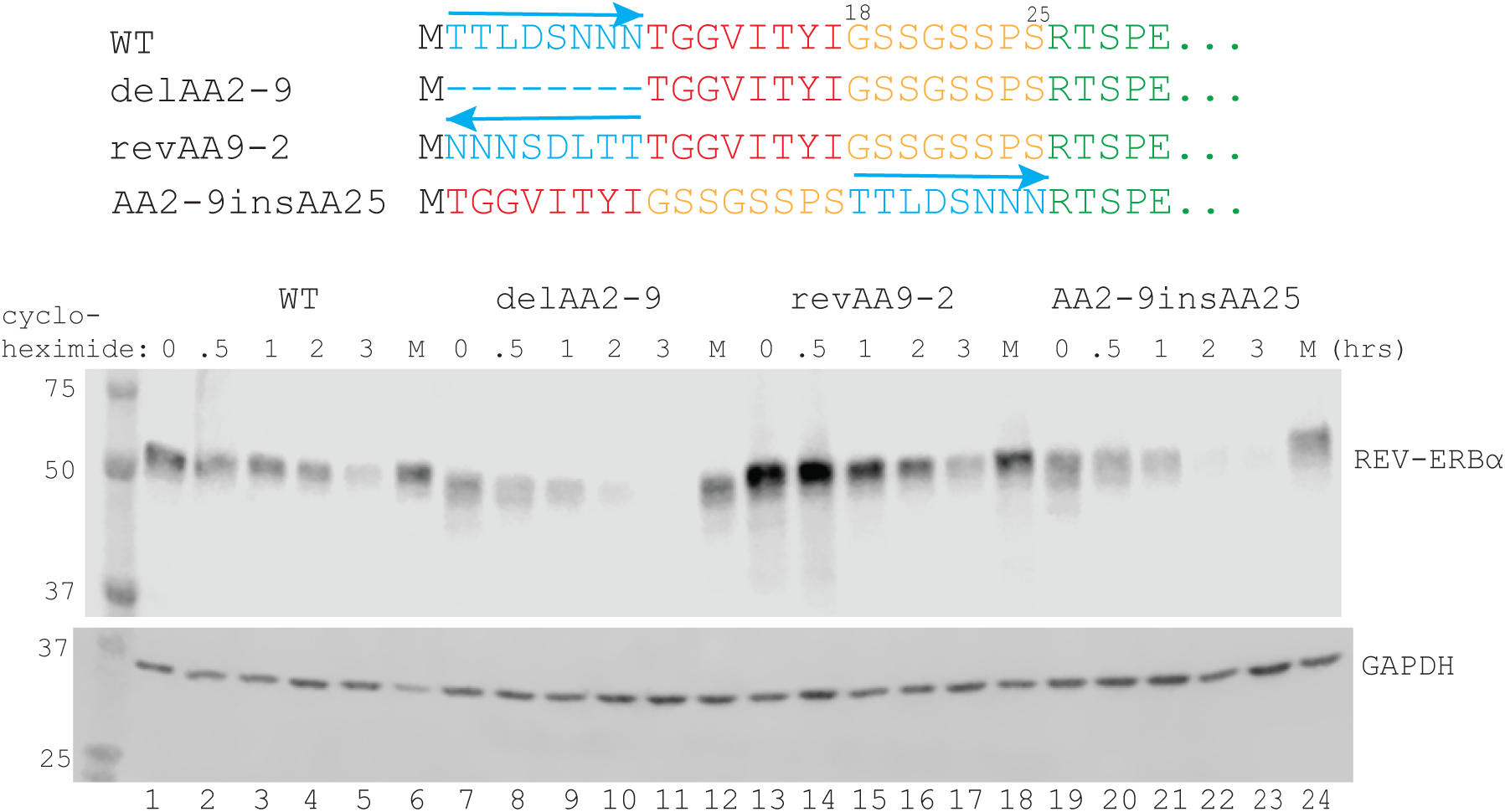
AA2-9 must be at the very N-terminus to protect REV-ERBα from fast degradation. AA2-9 was introduced back into the deletion mutant dAA2-9, but was now positioned as AA18-25, shown as mutant AA2-9insAA25. WT (lanes 1-6) or the indicated mutants (lanes 7-24) were transfected into HEK293 cells seeded in 24-well plate. 36-48 hours after transfection, cells were treated with cycloheximide for the indicated number of hours, and the Western blot was probed with anti-REV-ERBα antibody. The membrane was then probed with anti-glyceraldehyde-3-phosphate-dehydrogenase (GAPDH) antibody, as a loading control. Lanes “M” were lysates from cells treated with MG132 for 3 hours (lanes 6, 12, 18, and 24).

### AA4-9 is the site of interaction with the E3 ubiquitin-ligase SPSB4

Knocking out REV-ERBα has two major important effects on mice. First, these mice exhibit significantly shorter circadian period length to WT mice when kept at constant darkness. Second, mouse to mouse variation in the period length is much higher in the knockout mice when compared to the WT (4). These results suggested the loss of REV-ERBα led to a propensity to have an inaccurate clock.

Furthermore, we have demonstrated that when the level of the ubiquitin ligase SPSB4 was knocked down in U2OS cells, an increase in period length was observed, along with an apparent stabilization of REV-ERBα (16). These results imply that the site of interaction with SPSB4 on REV-ERBα could potentially control or regulate the circadian period. It is therefore important to locate the binding site of SPSB4 on REV-ERBα. The Drosophila homolog of SPSB protein, GUSTAVUS, was first shown to bind the Drosophila RNA helicase, VASA, (26). Human SPSB1, 2, and 4 were later shown to bind and degrade Par4 (26, 27). The peptide sequence DINNN was identified as the binding site for SPSB1, 2, and 4 on inducible nitric oxide synthase (iNOS) as well (28). To map the region on REV-ERBα that is required for its binding to SPSB4, we introduced MYC- or HA-epitope into the C-termini of SPSB4 and REV-ERBα. This is essential since there is no reliable anti-SPSB4 antibody, and some of the deletions introduced into REV-ERBα would inevitably remove the epitope that is the site of recognition by the commercially available anti-REV-ERBα antibody. A series of large truncation mutants that covers the entire 615 AAs of the protein were first prepared in REV-ERBα-MYC. SPSB4-HA was paired with various deletion mutants of REV-ERBα-MYC, in co-transfection experiments, followed by IP and western blots with anti-

HA, anti-MYC, or anti-REV-ERBα antibody, where appropriate. As summarized in Figure 5, positive interactions between the two proteins are indicated by the “+” signs to the right of the corresponding mutants. The first set of REV-ERBα-MYC deletion mutants (Figure 5A) helped narrow down the SPSB4-binding domain to the first 200 amino acids, as its removal from two mutants (201-400MYC and 401-615MYC) abolished their ability to interact with SPSB4-HA. The next set of REV-ERBα-HA mutants with smaller deletions were made and tested for binding to SPSB4-MYC, and the results narrowed down the binding to the first 50 amino acids (Figure 5B), as its removal from the two deletion mutants (51-400HA and 101-400HA) abolished binding to SPSB4.

**Figure 5.**
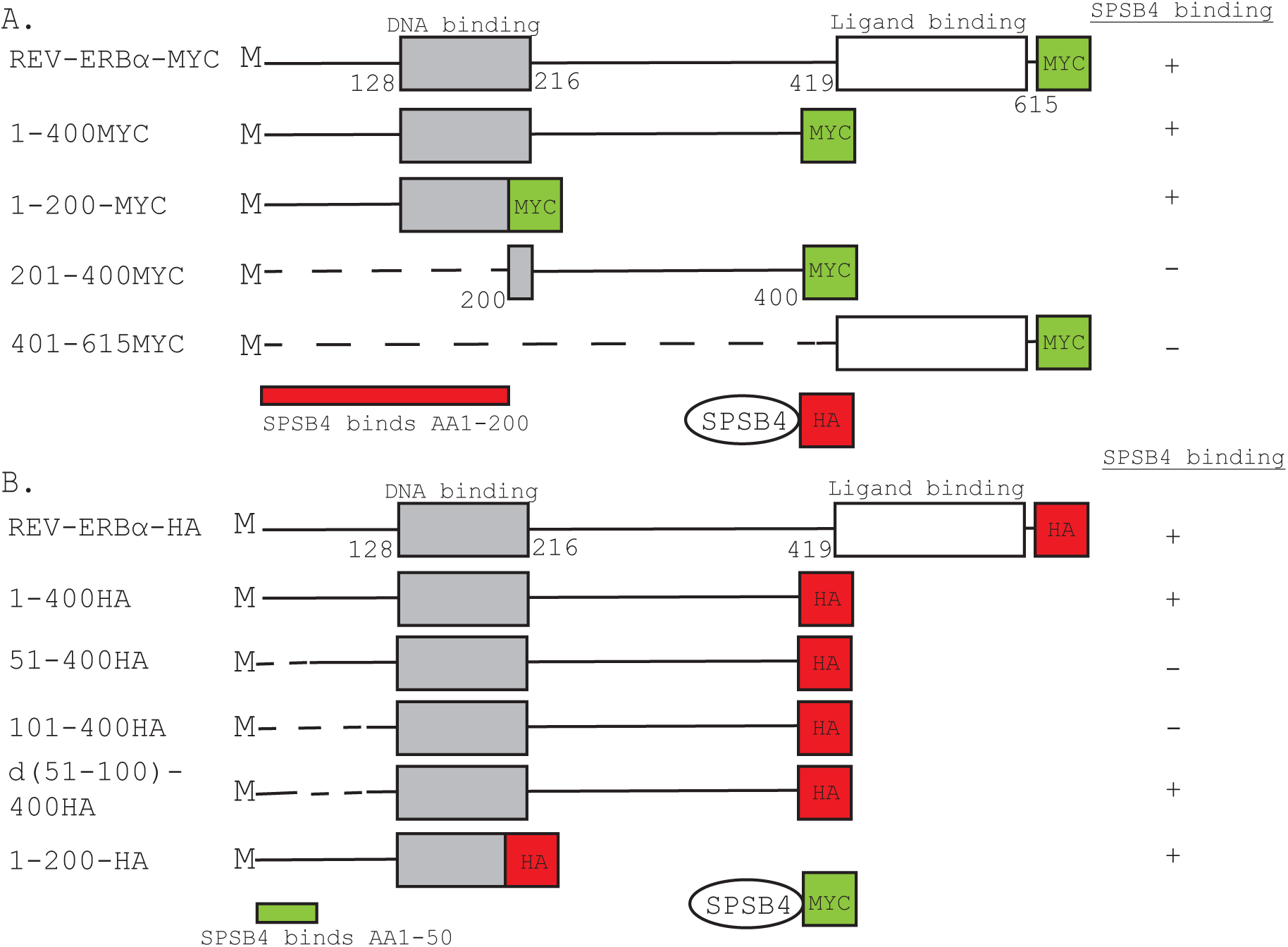
Localization of SPSB4 binding to the N-terminus AA1-50 of REV-ERBα. MYC- and HA-epitopes were inserted into the C-terminus of REV-ERBα and SPSB4, by in vitro mutagenesis. Various truncations and deletions were then introduced into REV-ERBα-MYC and REV-ERBα-HA to generate the deletion mutants as shown in 5A and 5B. A. WT REV-ERBα-MYC and large truncation mutants were co-transfected along with SPSB4-HA into HEK293. 36-48 hours after transfection, cell lysates were obtained and mixed with either anti-MYC or anti-HA antibody, followed by immunoprecipitation (IP) with Protein A agarose beads. Western blots were probed with anti-REV-ERBα, anti-MYC antibody, or anti-HA antibody, where appropriate. Binding to SPSB4 are shown as “+” or “-” sign to the right of each construct. The combined data narrowed down the binding of SPSB4 to the first two hundred amino acids of REV-ERBα (red bar, AA1-200). B. WT REV-ERBα-HA and the indicated deletion mutants were co-transfected with SPSB4-MYC into HEK293. 36-48 hours after transfection, cell lysates were obtained and mixed with either anti-HA or anti-MYC antibody, followed by immunoprecipitation (IP) with Protein A agarose beads. Western blots were probed with anti-REV-ERBα, anti-MYC antibody, or anti-HA antibody, where appropriate. Binding to SPSB4 are indicated by “+” sign to the right each construct. The combined data narrowed the site of interaction to the first fifty amino acids of REV-ERBα (green bar, AA1-50).

Next, we constructed additional N-terminal deletion mutants to further localize the binding site for SPSB4. As shown in Figure 6A, these are full length REV-ERBα protein with only the indicated length of peptide deleted (dashed-lines in the sequence). IP with anti-REV-ERBα antibody was successful in pulling down the WT and the indicated N-terminal mutants (lanes 5-8, around 75kD, marked by open arrowhead). Most importantly, although similar levels of SPSB4 were expressed (lanes 9-12, detected by anti-MYC antibody, around 25kD, marked by solid arrowhead), it was only brought down by the WT (lane 13) or the delAA10-25 mutant (lane 15) (also marked by the “+” sign to the right of the corresponding sequence). These data suggest that SPSB4 was bound to AA2-9 (blue sequence), as its absence in delAA2-17 and delAA2-25 abolished the binding (lanes 14 and 16, also indicated by the “-” sign to the right of the sequence).

**Figure 6.**
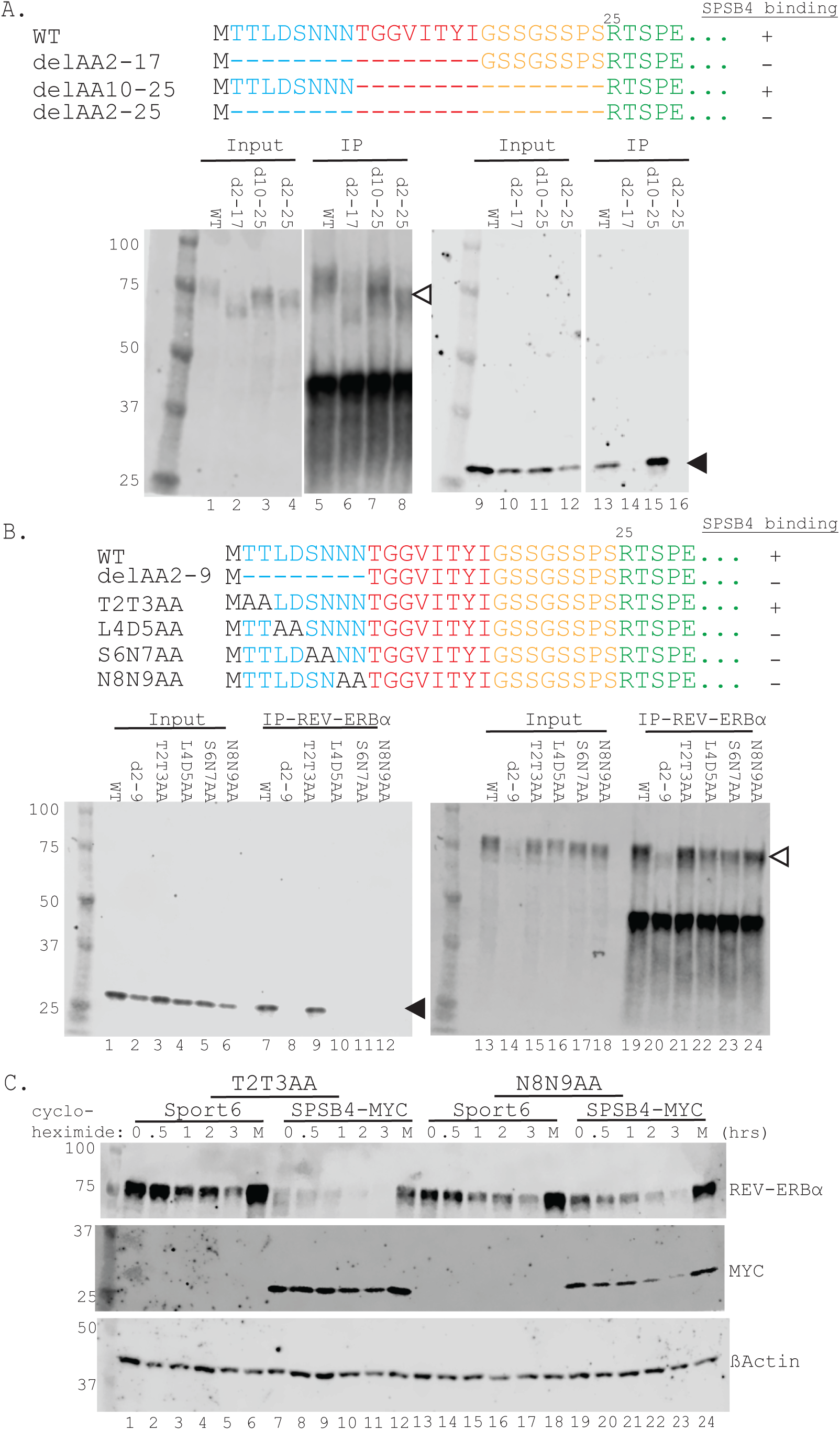
Localization of SPSB4 binding site to AA4-9 of REV-ERBα. A. WT REV-ERBα and the indicated mutants (lanes 1-4) were transfected along with SPSB4-MYC, and cells lysates were obtained 36-48 hours after transfection. Immunoprecipitation was carried out (lanes 5-8) with an anti-REV-ERBα antibody. The western blots were probed with anti-REV-ERBα (lanes 1-8) or anti-MYC antibody (lanes 9-16). The positions of REV-ERBα and SPSB4 are marked with open and solid arrowhead, respectively. Binding and non-binding mutants are indicated by “+” and “-” signs, respectively, shown to the right of the sequence. The combined data narrowed down the binding of SPSB4 to AA2-9. B. Double alanine substitution mutants of REV-ERBα, used previously in Figure 2, were tested in binding assays with SPSB4-MYC, as performed in 6A. Transfected cell lysates were IP with anti-REV-ERBα antibody (lanes 7-12 and lanes 19-24). The western blots were probed with anti-MYC antibody (lanes 1-12) or anti-REV-ERBα antibody (lanes 13-24). The positions of REV-ERBα and SPSB4 are marked with open and solid arrowhead, respectively. C. The two mutants used in Figure 6B, T2T3AA (lanes 1-12) and N8N9AA (lanes 13-24) were transfected into HEK293 cells along with either Sport6 or SPSB4-MYC, as indicated. 36-48 hours after transfection, cells were treated with cycloheximide for the indicated number of hours, and the Western blot was probed with anti-REV-ERBα antibody. The membrane was then probed with anti-glyceraldehyde-3-phosphate-dehydrogenase (GAPDH) antibody, as a loading control. Lanes “M” were lysates from cells treated with MG132 for 3 hours (lanes 6, 12, 18, and 24). Expression of SPSB4 was detected by anti-MYC antibody, since the SPSB4 was tagged with a MYC-epitope at its C-terminus.

**Figure 7.**
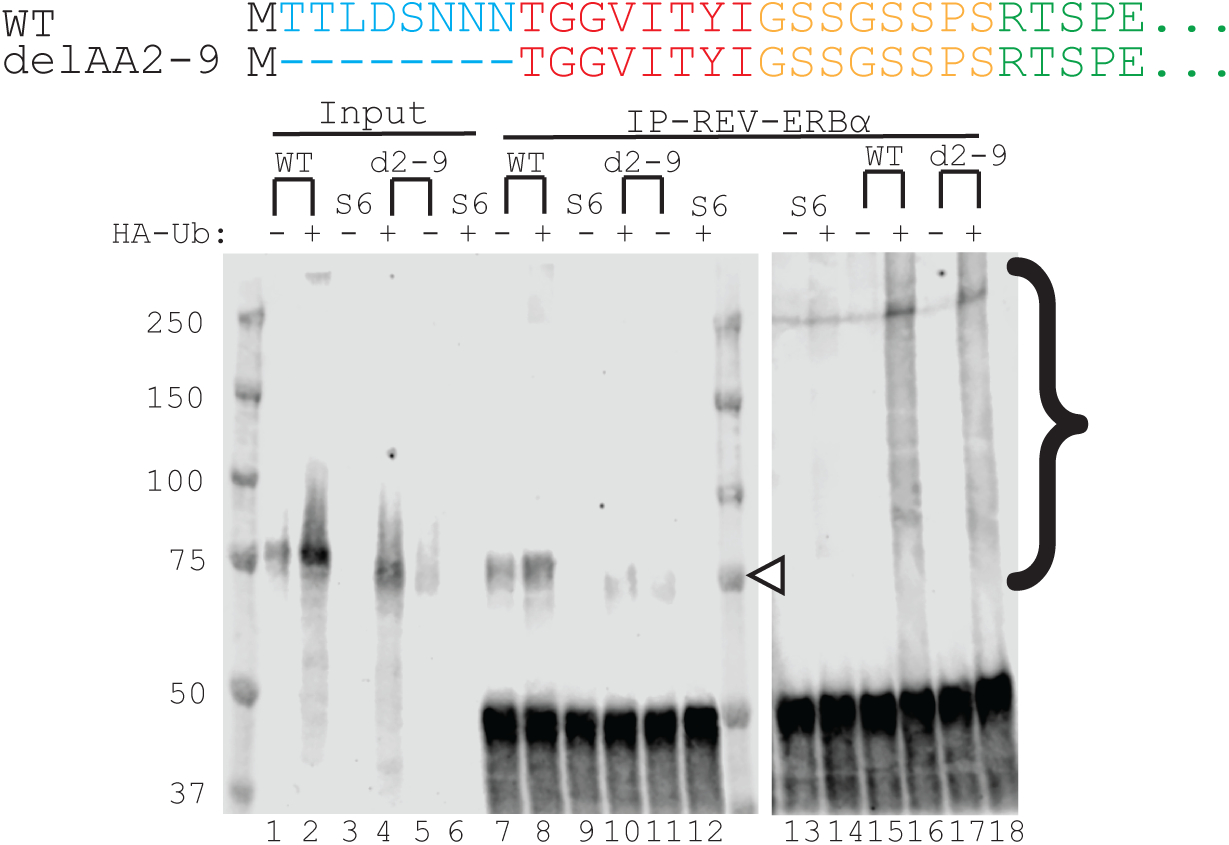
Deleting AA2-9, the binding site of SPSB4, did not have much effect on ubiquitination of REV-ERBα. Sport6, WT, or delAA2-9 were transfected into HEK293 cells with (“+” lanes) or without (“-” lanes) co-transfection of a plasmid that expresses a HA-tagged ubiquitin (HA-Ub). Lysates from transfected cells (lanes 1-6) were immunoprecipitated (IP) with an anti-REV-ERBα antibody (lanes 7-18). Western blots were probed with an anti-REV-ERBα (lanes 1-12) or anti-HA antibody (lanes 13-18). Bands that form a ladder are marked with a bracket on the right, representing REV-ERBα proteins that have increasing number of ubiquitin attached.

We then made use of the full-length REV-ERBα mutants with only two amino acids substitution (used in earlier experiment presented in Figure 3A) to determine the sequence required for SPSB4 binding. As shown in Figure 6B, WT REV-ERBα and all the mutants were brought down by IP with anti-REV-ERBα antibody (lanes 19-24, marked by open arrowhead). Despite the detection of SPSB4-MYC in all the input samples (lanes 1-6, marked by solid arrowhead) and, consistent with the results in Figure 6A, binding to the WT protein was clearly detected (lane 7), but was abolished with delAA2-9 (lane 8). Out of the four substitution mutants (lanes 9-12), only T2T3AA was still able to bind with SPSB4 (lane 9, and indicated by the “+” sign to the right of the sequence). Importantly, T2T3AA was still highly susceptible to degradation induced by SPSB4 (Figure 6C, compare lanes 7-12 to lanes 1-6). Consistent with the loss of binding to SPSB4 (Figure 6B), N8N9AA became more resistant to degradation by SPSB4 (Figure 6C, compare lanes 19-24 to lanes 13-18). The combined data show that SPSB4 binding site is the six amino acid peptide LDSNNN at AA-4-9 of REV-ERBα.

### Does AA2-9 affect ubiquitination of REV-ERBα stability?

Data shown in Figures 5 and 6 clearly demonstrated that AA4-9 is the binding site for the ubiquitin ligase SPSB4, but the removal of AA2-9 resulted in faster degradation (Figure 2), which is opposite to what expected when an ubiquitin ligase binding site is removed. Furthermore, the efficient ubiquitination of K20R (Figure 1D) implicates possible N-terminal ubiquitination, it is therefore important to test if and how ubiquitination is affected when AA2-9 is deleted. Both WT and delAA2-9 were transfected into HEK293 cells, with (Figure 8, “+” lanes) or without (“-” lanes) co-transfection of a plasmid that expresses HA-tagged ubiquitin cDNA (HA-Ub). Cells were lysed and followed by immunoprecipitation (IP) with anti-REV-ERBα antibody (lanes 7-18). Western blot using anti-REV-ERBα antibody showed that both WT (lanes 7,8) and delAA2-9 (lanes 10,11) were successfully brought down by IP (position marked by open arrowhead at 75kD). Consistent with delAA2-9 being less stable, higher levels of WT were detected both in the input (compare lanes 1 and 2 to lanes 4 and 5), and the IP samples (compare lanes 7 and 8 to lanes 10 and 11). Meanwhile, endogenous REV-ERBα was undetectable in HEK293 cells transfected with the empty vector Sport6 (“S6” lanes 3, 6, 9, and 12). When anti-HA antibody was used in the western blot (lanes 13-18), bands appeared as ladders (marked by bracket) were detected only in those lanes where HA-Ub were also co-transfected (lanes 16 and 18). These bands all have higher molecular weight than that of REV-ERBα, suggesting that they are REV-ERBα with increasing numbers of ubiquitin molecules attached, as each additional ubiquitin molecule would increase the size of the protein by 8kD. The absence of these ladders in the HEK293 cells transfected with Sport6 (lanes 13 and 14) further confirmed their specificity. Therefore, although there was less delAA2-9 immunoprecipitated than the WT protein (compare lane 10 to lane 8), the ladder of bands detected in delAA2-9 (lane 18) were very similar to those detected with the WT (lane 16). It is difficult to make accurate quantitation from an immunoprecipitation experiment, but the results suggested a higher tendency for delAA2-9 to be ubiquitinated, consistent with this mutant being more unstable.

**Figure 8.**
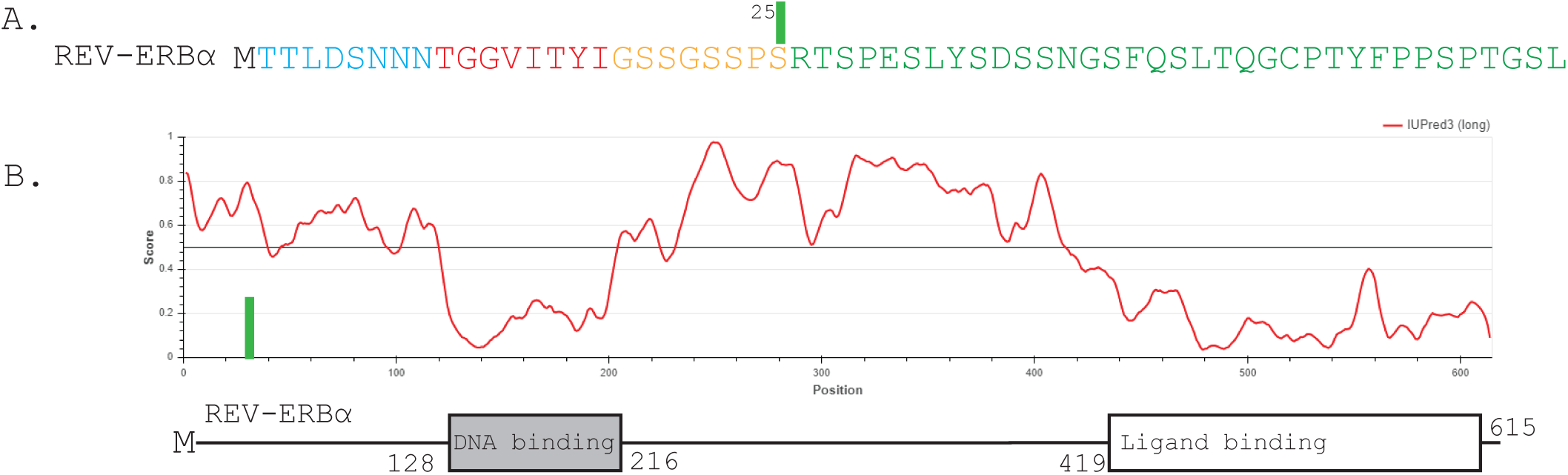
The N-terminus of REV-ERBα is predicted to be an intrinsic disordered region. A. The first 50 amino acids of REV-ERBα is shown, with the same color scheme as used in this study. B. Profile of predicted disordered regions across the full 615 amino acids of REV-ERBα (bottom). The green bar in 8B matches marks the position of the first 25 amino acids in 8A. All scores over 0.5 on the Y-axis predicts an intrinsic disordered region. Protein sequences of REV-ERBα was input into https://iupred3.elte.hu (37) to obtain the profile.

## Discussions

We and others have investigated how various ubiquitin ligases contribute to the degradation of REV-ERBα, and their effects on the circadian clock (15–19). In this study, we focused on trying to understand the central events that need to happen to REV-ERBα protein itself for degradation to take place. As ubiquitin ligases drive degradation by initiating the whole process through ubiquitination of the target protein on certain lysine residue(s), we were surprised by the results that lysines are not required for ubiquitination and degradation of REV-ERBα (Figure 1). A limited number of proteins have been described to undergo lysine-independent ubiquitination and degradation, leading to the discovery of N-terminual ubiqutination and degradation (21–25). The efficient ubiquitination and degradation of the K20R mutant (Figure 1) suggested that REV-ERBα may be added to this short list of proteins (25). One caveat of these reported N-terminal ubiquitinated proteins is that they could only be demonstrated by using an *in vitro* translation system, where the free α-amino group of the N-terminal methionine could be modified and blocked from potential ubiquitination in a cell-free ubiquitination assay. There is no easy way to modify the N-terminus of a freshly translated protein *in vivo* without affecting its expression. This is further complicated by possible effects mediated by the N-end rule of degradation (29), which recently was renamed as N-degron rule (30). These technical difficulties may be reasons why there are not many follow-up studies or other proteins reported to use such N-terminal ubiquitination mechanism. In recent years, there have been renewed interests in lysine-independent ubiquitination and their effects, many of which however, are not related to degradation (31).

It was intriguing to see the big change in stability between delAA2-9 and delAA2-17 (Figure 2), but these results provided a first glimpse into how this region may control the stability of REV-ERBα. When AA2-9 was deleted, the protein became unstable (Figure 2), suggesting that it plays an important role in maintaining the native protein in a relatively stable (WT) conformation, such that REV-ERBα can carry out its functions during times of the day when it is needed. The similarly unstable re-aligned mutant (AA2-9insAA25, Figure 4) indicated that AA2-9 must be located at the very N-terminus, for its degradation-impeding function.

The apparent lack of specific sequence requirement for AA2-9 to maintain the WT-like stability (Figure 3) may appear puzzling. However, others have described proteins that were targeted through an ubiquitin-independent proteasomal degradation pathway, mostly attributed to an intrinsic disordered region (32–35).

Intrinsic disordered region has gained a lot of interest in recent years (36), as it has the flexibility to assume many different structures, through post-translational modification, or various intra- and inter-molecular interactions. When the REV-ERBα protein sequence was subjected to a context-dependent prediction for protein disorder (37), the first 100 AAs are predicted to be largely disordered (Figure 8B, the higher the Score is above 0.5 on the Y-axis, the higher possibility of being disordered). This may explain why small substitution and even altering the orders without changing the composition of AA residues within AA2-9 did not affect the protein stability (Figure 3A-E). On the other hand, smaller deletions, delAA2-3 and AA2-7 resulted in slightly faster degradation relative to the WT protein (Figure 3G and 3F), but not to the extent of delAA2-9, suggesting that the degradation-protecting function is length dependent. AA2-9 is perhaps optimal for the protective effect, while shortened length still maintains a weakened but discernable activity. This in a way is consistent with the possible N-terminal ubiquitination, although we did not have direct evidence.

At first sight, the mapping of the SPSB4 binding site to AA4-9 is counterintuitive, in that removal of a E3-binding motif should result in a more stable protein. This contradicts the fact that deleting AA2-9 from the WT resulted in a faster degradation (Figure 2). One possible explanation for these apparently contradictory results is that AA2-9 serves two separate functions. First, it is the binding site to recruit SPSB4, thus initiating the first step of ubiquitination to mark and prime the protein for its subsequent interaction with the 26S proteasome. Second, AA2-9 at the N-terminus may function to mask and impede a peptide sequence that is a potential proteasome initiation region, as described in some proteins (38–42), which when exposed, will result in one or more of the following events, accelerated interaction with the proteasome, unfolding of the protein, and translocation into the pore of catalytic core of the proteasome (13,14), where the actual degradation takes place (42,43). Weakened masking effects may also explain the weaker but discernable degradation-protective effects observed with the smaller deletions (delAA2-3, delAA2-7) as compared to delAA2-9 (Figures 3F and 3G). A potential site of such a proteasome initiation region on REV-ERBα may be AA10-17, which cloud explain why the delAA2-17 mutant became a very stable protein and even MG132 resistant (Figure 2), as both the SPSB4-binding capacity (AA2-9) and the proteasome initiation potential have been removed. The stability of the re-positioned mutant (AA2-9insAA25, Figure 4) being like delAA2-9 suggested that the realignment of this region may have disrupted how different elements within the N-terminus interact with each other, thus affecting the overall stability of REV-ERBα. The stability of additional mutants in AA10-25 are now being studied to further understand its involvement in regulating the degradation process.

UBE2W is the first E3 ligase shown to promote N-terminal ubiquitination (44–46). A recent study identified 73 putative substrates of UBE2W (47), most of which are predicted to have disordered N-termini. Among these were UCHL1 and UCHL5, two related deubiquitinases, that were identified as targets of N-terminal ubiquitination (47). Our mapping of the SPSB-4 binding site to the N-terminus of REV-ERBα (Figures 5 and 6), a region predicted to be intrinsically disordered (Figure 8), and the possible N-terminal ubiquitination, are features found on some substrates of UBE2W (45–47). These similarities in some ways validate the importance of this N-terminal degradation control region in regulating REV-ERBα stability. Nevertheless, a major difference must be noted in that UBE2W only conjugates a single ubiquitin (mono-ubiquitination) onto its substrate (46,47), and N-terminal ubiquitination by UBE2W is not a strong signal for degradation, instead it seems to modulate the deubiquitinase function of those two substrates (47). In contrast, REV-ERBα was clearly poly-ubiquitinated and degraded (Figure 1).

In conclusion, we have several interesting findings on the mechanisms of regulating REV-ERBα degradation. This includes lysine-independent ubiquitination and degradation, which implies perhaps N-terminal ubiquitination. The properties of these regulatory elements fit into a general theme of how degradation is controlled for many different proteins (48–51), but they appear to be all packaged into the first 25AAs at the N-terminus for REV-ERBα. The degradation blocking activity of AA2-9 appears to be non-sequence specific but length dependent, explainable by the composition of these amino acids that are predicted to be an intrinsic disordered region. Another function of AA2-9 is its ability to bind the ubiquitin ligase SPSB4, this is exciting since our previous experiments showed that SPSB4-knockdown will result in the stabilization of REV-ERBα, as well as a longer circadian period (16). An critical shortfall of these ubiquitin ligase knockdown experiments is that one can always argue that the observed effects may be due to other possible targets of the ubiquitin ligase. Now, we can directly mutate the specific binding site on REV-ERBα and observe its effect on the circadian period, and in the future, even in animal’s behavior and metabolism. Overall, the discovery of this degradation-control domain opens up the possibility to study how REV-ERBα existence in the cells is only limited to a specific time interval in the day (ZT6 to ZT10).

Recently, Cryo-EM was able to resolve the detailed interactions (52–53) of various components with the proteasome throughout the degradation process. Similar approach would help the understanding of the structural changes that may be the results of various possible interactions among different segments within the N-terminus, as well as its binding to SPSB4, and how these interactions actually control the degradation process.

## Acknowledgement

This work is supported by NIH NIGMS grants GM109861 and GM127044 to Jason P. DeBruyne, RCMI grant U54MD007602 to Ting-Chung Suen, as well as in part by NIH NINDS grant U54 NS083932 and NIH NIMHD grants 8G12MD007602, 8U54MD00758, 1G20RR031196, S21MD000101, C06RR18386 to Morehouse School of Medicine.

## Disclaimer

The content is solely the responsibility of the authors and does not necessarily represent the official view of the National Institute of Health.

## Conflict of Interest

The authors declare that they have no conflicts of interest with the contents of this article.

## Notes

### Competing Interest Statement

The authors have declared no competing interest.

### Summary of Updates

We have corrected mistakes and made changes for the manuscript to be more easily understandable. We have shortened the discussions and revised Figure 8 to reduce speculation.

